# CDC6 regulates mitotic CDK1 via cyclin B and not cyclin A and acts through a *bona fide* CDK inhibitor Xic1

**DOI:** 10.1101/2020.06.13.149989

**Authors:** Mohammed El Dika, Lisa Wechselberger, Bilal Djeghout, Djamel Eddine Benouareth, Krystyna Jęderka, Sławomir Lewicki, Claude Prigent, Malgorzata Kloc, Jacek Z. Kubiak

## Abstract

The timing of the M-phase entry and its progression are precisely controlled by a CDC6-dependent mechanism that inhibits the major mitotic kinase CDK1, and, thus, regulates the dynamic of CDK1 during the M-phase. In this paper, we describe the differential regulation of the mitotic CDK1 dynamics by exogenous cyclin A or a non-degradable cyclin B added to the *Xenopus laevis* embryo cycling extracts. We showed that the variations in the level of cyclin B modify both CDK1 activity and the timing of the M-phase progression, while the cyclin A levels modify only CDK1 activity without changing the timing of the M-phase events. In consequence, CDC6 regulates the M-phase through endogenous cyclin B, but not cyclin A, which we demonstrated directly by the depletion of cyclin A, and the addition of CDC6 to the cycling extracts. Further, we showed, by p9 precipitation (p9 protein associates with Cyclin-Dependent Kinases, CDK), followed by the Western blotting that CDC6, and the *bona fide* CDK1 inhibitor Xic1, associate with CDK1 and/or another CDK present in Xenopus embryos, the CDK2. Finally, we demonstrated that the Xic1 temoprarily separates from the mitotic CDK complexes during the peak of CDK1 activity. These data show the differential coordination of the M-phase progression by CDK1/cyclin A and CDK1/cyclin B, confirm the critical role of the CDC6-dependent CDK1 inhibition in this process and show that CDC6 acts through the cyclin B- and not cyclin A/CDK complexes. This CDC6- and cyclin B-dependent mechanism may also depend on the precisely regulated association of Xic1 with the CDK complexes. We postulate that the dissociation of Xic1 from the CDK complexes allows the maximal activation of CDK1 during the M-phase.

## Introduction

The mitotic cell cycle is composed of four phases: G1, S, G2, and M, during which the eukaryotic cell produces two diploid, genetically identical daughter cells. The main two families of proteins involved in cell cycle control are cyclin-dependent kinases (CDKs) and cyclins. Several CDKs have been identified as being active during the cell cycle (Walker and Maller, 1991). The main CDKs that act during interphase are CDK2, CDK4, and CDK6, while the CDK1 is essential for the M-phase.

Cyclins, the regulatory subunits of CDKs, are divided into two categories, the G1/S cyclins and mitotic cyclins (D’Angiolella et al., 2001; Girard et al., 1991; Müller-Tidow et al., 2004). The G1/S cyclins bind to CDK2 and CDK4-6 during the G-phase and are necessary for the entry to and progression of the S-phase. The mitotic cyclins, which associate with CDK1 and CDK2 during the G2 phase, are required for entry into M-phase (Walker and Maller, 1991). The levels of mitotic cyclins A and B fluctuate during the cell cycle in contrast to CDK1 level, which remains stable during the whole cell cycle. The increase or decrease in cyclins’ level allows the cycling activation of CDKs with which they are associated (Evans et al., 1983; Pines, 1995; Pines and Hunter, 1991). The activity of CDKs is also regulated by the CDK inhibitors called CKIs. The CKIs bind to CDKs alone or CDK/cyclin complex and down-regulate the kinase activity. Two distinct classes of CDK inhibitors have been identified: INK4 and Cip/Kip (Blain et al., 1997; Sherr and Roberts, 1999). In *Xenopus laevis*, the Cip/Kip-type of CKI named Xic1, inhibits DNA replication through the binding to CDK2-cyclins complexes (You et al., 2002). So far, there are no data available on the involvement of the CKI in the mitotic CDK1 regulation in *Xenopus laevis* embryos or oocytes, however, such regulation was described in yeasts (Örd et al., 2019).

Cyclins are coded by at least 15 different genes in the human genome (Gopinathan et al., 2011). Only some of them are expressed in *Xenopus laevis* oocytes and embryos. B-type cyclins are major regulators of CDK1 during the M-phase (Arellano and Moreno, 1997; King et al., 1994). A-type cyclins also interact with CDK1 promoting entry into M-phase (Lorca et al., 1992; Pagano et al., 1992). Both cyclins A and B contain a destruction box required for their proteolysis via ubiquitination during the pertinent period of the cell cycle (Glotzer et al., 1991; Rechsteiner and Rogers, 1996). At prometaphase-metaphase transition, the ubiquitin ligase APC/C^Cdc20^ complex becomes activated inducing degradation of cyclin B and securin, which is necessary for the transition from the metaphase to the anaphase.

The cyclins are not the only key factors driving the embryonic cell cycle and determining the timing of the M-phase. There is another important factor involved in the process, which is the Cell Division Cycle 6, CDC6. We showed previously that this protein, by inhibiting CDK1, determines the timing of M-phase entry and progression in *Xenopus laevis* cell-free extract, and mouse embryo (Borsuk et al., 2017; El Dika et al., 2014a).

The CDC6 protein is an evolutionarily conserved member of the AAA + ATPase family that plays a key role in many cell functions, such as folding, unfolding, and degradation of proteins, vesicular transport, and the assembly of macromolecules for DNA replication (Duderstadt and Berger, 2008). The CDC6 contains the 200-250 amino acid long ATPase domain. The N-terminal region of CDC6 has one phosphorylation site for PLK1 and three consensus sites for phosphorylation by CDKs. These sites are phosphorylated in S-phase. It also has a leucine zipper domain, which is a domain of interaction with other proteins, and a so-called cyclin binding domain, which allows association with cyclins (Örd et al., 2019). This cyclin-binding motif is present in different substrates and inhibitors of CDK2 (Chen et al., 1996; Zhu et al., 1995), and binds specifically to the hydrophobic region of cyclin A (Russo et al., 1996). This improves the interaction between CDK2 and its targets. The D-boxes and Ken-boxes, present in CDC6 are necessary for ubiquitination by APC/C^Cdh1^, which induces CDC6 degradation by the proteasome during the G1/G0 phase (Petersen et al., 2000). During the M-phase, in HeLa cells, the CDC6 is phosphorylated by Polo-like kinase 1 (Plk1) (Yim and Erikson, 2010). CDC6 associates with Plk1 and localizes to the mitotic spindle throughout the metaphase and anaphase. The high CDC6 phosphorylation level correlates with the high Plk1 activity, and conversely, CDC6 is dephosphorylated in cells depleted of Plk1. Phosphorylation of CDC6 by Plk1 is required for the efficient interaction with CDK1 and for its inhibition. In yeast, CDC6 homolog has been shown to regulate CDK1 inactivation during the M-phase exit. The N-terminal region of CDC6 interacts with CDK1/cyclin B during M-phase exit and inhibits histone H1 kinase activity in vitro (Elsasser et al., 1996). The deletion of the CDK1 interaction domain in CDC6 slows down the M-phase exit (Calzada et al., 2001).

It has been shown recently in *Xenopus laevis* embryos that cyclin A is involved in the initiation of the M-phase by triggering the activation of CDK1/cyclin B complex (Vigneron et al., 2018). The question is whether CDC6 regulates CDK1/Cyclin A during M-phase in *Xenopus laevis* embryos. We showed that CDC6 is involved in the regulation of CDK1/Cyclin B in Xenopus, and mouse one-cell embryos (El Dika et al., 2014a). Here we demonstrate the presence of the regulatory network between cyclin A, cyclin B, CDK1 (and plausibly CDK2), and CDC6, and a potential role of CDK1/2 *bona fide* inhibitor Xic1 in the progression of the M-phase. This regulatory mechanism is of special importance for the coordinated activation of the CDK1 and the control of its amplitude during early embryo cleavage divisions when the precise timing of cell cycle events is required for the coordination with the genetic developmental program of the embryo.

## Material and methods

### Xenopus egg collection and activation

*Xenopus laevis* eggs were collected from the overnight spawning, dejellied with 2% L-cysteine pH 7.81 in XB buffer (100 mM KCl, 1 mM MgCl2, 50 mM CaCl2, 10 mM HEPES, and 50 mM sucrose pH 7.6), washed in XB, activated with calcium ionophore A23187 at 0.5 mg/ ml, and then washed in XB as described before (Dika et al., 2014).

#### Xenopus cell free extracts

Cycling cell-free extracts were obtained from calcium ionophore-activated one-cell embryos as we described previously (El Dika et al., 2014a,b) using the method allowing biochemical analysis prior to the M-phase entry (see figure 1 in the results section).

**Fig.1.**
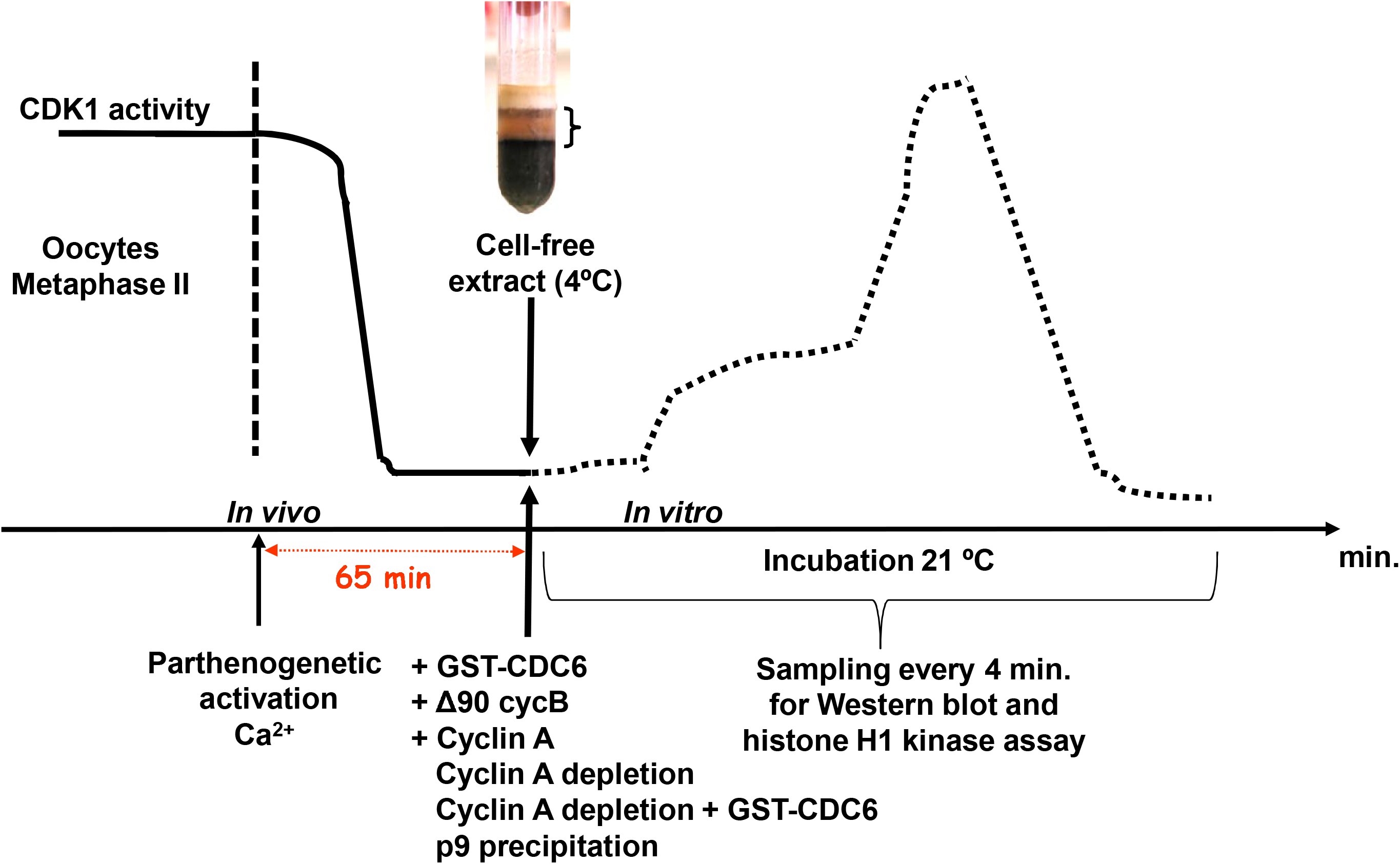
Outline of the experiments.

#### Anti-XlCDC6 production

Complementary DNA encoding wild-type *X. laevis* CDC6, recombinant protein, and the corresponding polyclonal antibodies were produced as previously described (El Dika et al., 2014a).

#### Cyclin A immunodepletion

Immunodepletion of Cyclin A from Xenopus embryo extracts was carried out using AffiPrep Protein A beads (Sigma) conjugated with the *Xenopus laevis* anti-Cyclin A1 (a gift from Daniel Fisher, IGMM, Montpellier France) or with the pre-immune serum, overnight at 4°C. 200ml of beads were washed four times with XB buffer (pH 7.6) and incubated with 400 ml of the premitotic extracts. Following 30 min incubation at 4°C, extracts were spun down, beads were removed, and the supernatant was recovered. Two runs of the immunodepletion were necessary to remove 90% of Cyclin A from the extract.

#### Immunoblotting of *X. laevis* proteins

Proteins in the extracts were separated by SDS-PAGE (8 to 12.5% gels), transferred to the nitrocellulose membranes (Hybond C, Amersham Biosciences), and probed with primary antibodies against cyclin A1 (gift from Daniel Fisher, IGMM, Montpellier France), cyclin B2 and CDC27 (gift from Thierry Lorca, CRBM, Montpellier, France), MCM4 (gift from Marcel Méchali, IGH, Montpellier, France), Xic1 (gift from Anna Philpott, University of Cambridge, UK), CDC6 (produced by us), and PSTAIR (Sigma). The secondary antibodies were: alkaline phosphatase-conjugated anti-rabbit or anti-mouse IgG. Bands were visualized with Enhanced Chemifluorescence reagent (ECF; Amersham Biosciences) and quantified with ImageQuant 5.2 software (Amersham Biosciences).

#### Recombinant proteins added to extracts

Purified human cyclin A (a gift from Daniel Fisher, IGMM, Montpellier, France) or non-degradable Δ90 cyclin B from sea urchin (a gift from Marie-Anne Felix, IJM, Paris, France) were added to the extracts.

#### Sepharose p9 beads precipitation

The p9-Sepharose beads (a gift from L. Meijer and O. Lozach from the Marine Station, Roscoff, France) were used to precipitate CDK complexes (CDK1 and CDK2 in *X. laevis* embryo). The 10 μl of extracts were mixed with 10 μl p9 beads pre-equilibrated with the homogenization buffer (MOPS pH 7.2, 60 mM β-glycerophosphate, 15 mM EGTA, 15 mM MgCl2, 2 mM dithiothreitol, 1 mM sodium fluoride, 1 mM sodium orthovanadate and 1 mM disodium phenyl phosphate) containing 1% BSA, 1mM AEBSF, aprotinin, leupeptin, pepstatin, and chymostatin (10 μg/ml each). Subsequently, the samples were gently agitated for 2.5 h at 4°C. Following the first brief spin down (5,000 g, 1 min, 4°C), the supernatant was collected for the Western blot analysis, while the p9 beads pellet was washed four times with 1 ml of washing buffer (50 mM Tris-HCl pH 7.4, 250 mM NaCl, 5 mM EDTA, 5 mM EGTA, 5mM sodium fluoride and 0.1% Nonidet-P 40) supplemented with 0.5 mM AEBSF, aprotinin, leupeptin, pepstatin, and chymostatin (10 μg/ml each). Finally, the beads were resuspended in 12 μl of 2× Laemmli sample buffer and heated at 85°C for 5 min.

#### Histone H1 kinase activity assay

MPF activity was measured as we previously described (Dika et al., 2014a,b;). In short, samples of extracts were diluted in the MPF buffer supplemented with 0.5 mM sodium orthovanadate and protease inhibitors, and containing 0.4 mg/ml H1 histone (type III-S), 1 μCi [γ32P] ATP (specific activity: 3000 Ci/mmol; Amersham Biosciences) and 0.8 mM ATP. They were incubated at 30°C for 30 minutes, mixed with Laemmli sample buffer, and heated at 85°C. Following the SDS-PAGE, the histone H1-incorporated radioactivity was measured with STORM phosphorimager (Amersham Biosciences). Data were analyzed with ImageQuant 5.2 software.

### Results

#### The exogenous Cyclin A and cyclin B differentially regulate M-phase progression in *X. laevis* embryo cycling extract

To investigate the effects of cyclin A and cyclin B on the M-phase timing, we first verified when the cytoplasmic extracts, prepared from the parthenogenetic embryos, enter the M-phase. There were 3 groups of extracts: 1. Control, without the addition of any purified protein, 2. with the addition of three concentrations of purified human cyclin A (a gift from Daniel Fisher, IGMM, Montpellier, France) 3. With the addition of non-degradable Δ90 cyclin B from sea urchin (a gift from Marie-Anne Felix, IJM, Paris, France). The prepared extracts were then incubated at 21□C, and samples were collected every 4 min for Western blot analysis with primary antibodies against CDC27, and histone H1 kinase assay. The CDC27, a subunit of the APC/C, is a direct substrate for CDK1 phosphorylation. On Western blot, the phosphorylation of CDC27 is visible as the up-shifted band resulted from the phosphorylation by active CDK1 (Huang et al., 2007). Figure 2A shows the timing of the CDC27 up-shift in extracts with cyclin A addition. Figure 2B shows the timing of the CDC27 up-shift in the extract with cyclin B addition.

**Fig. 2.**
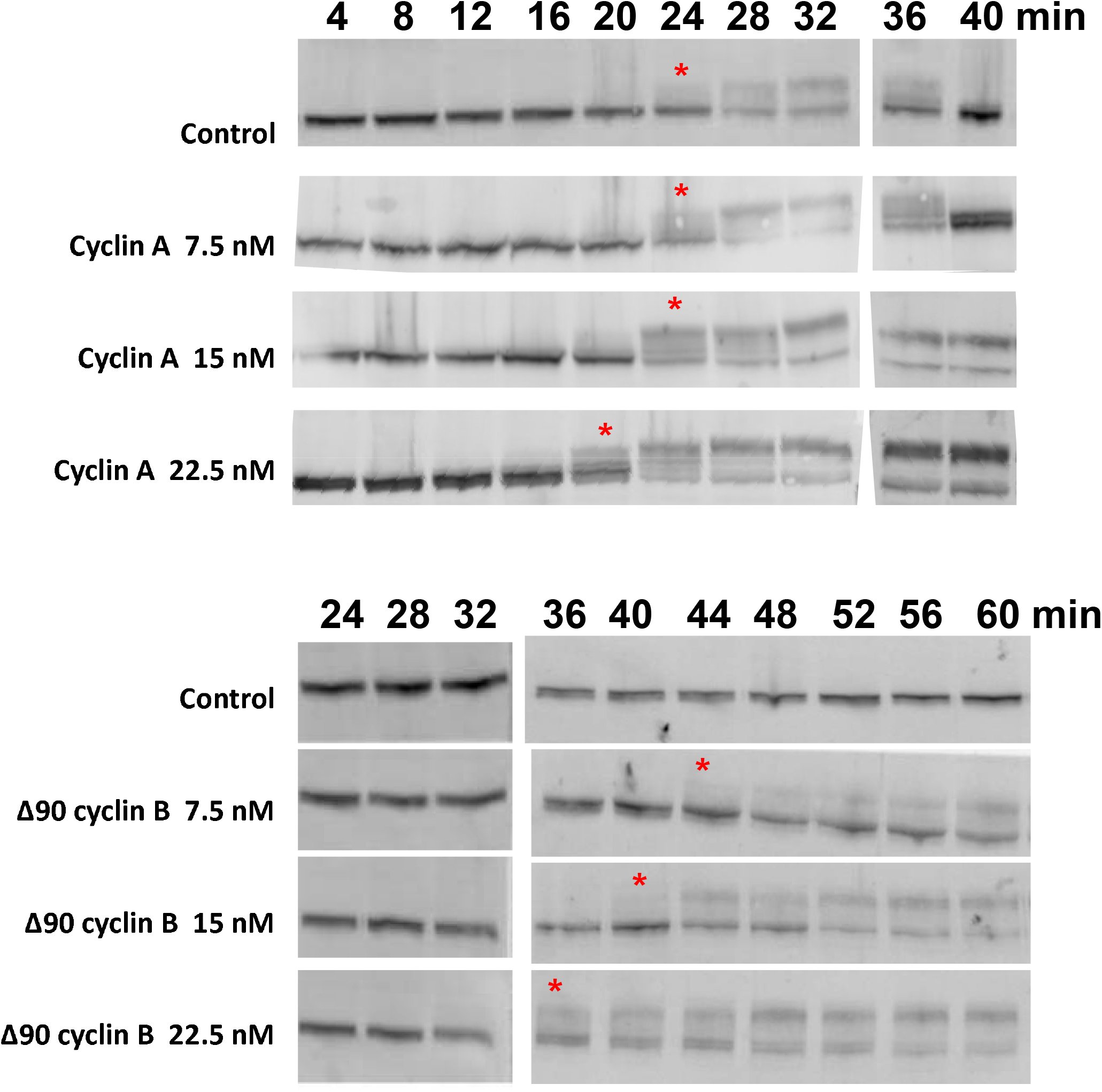
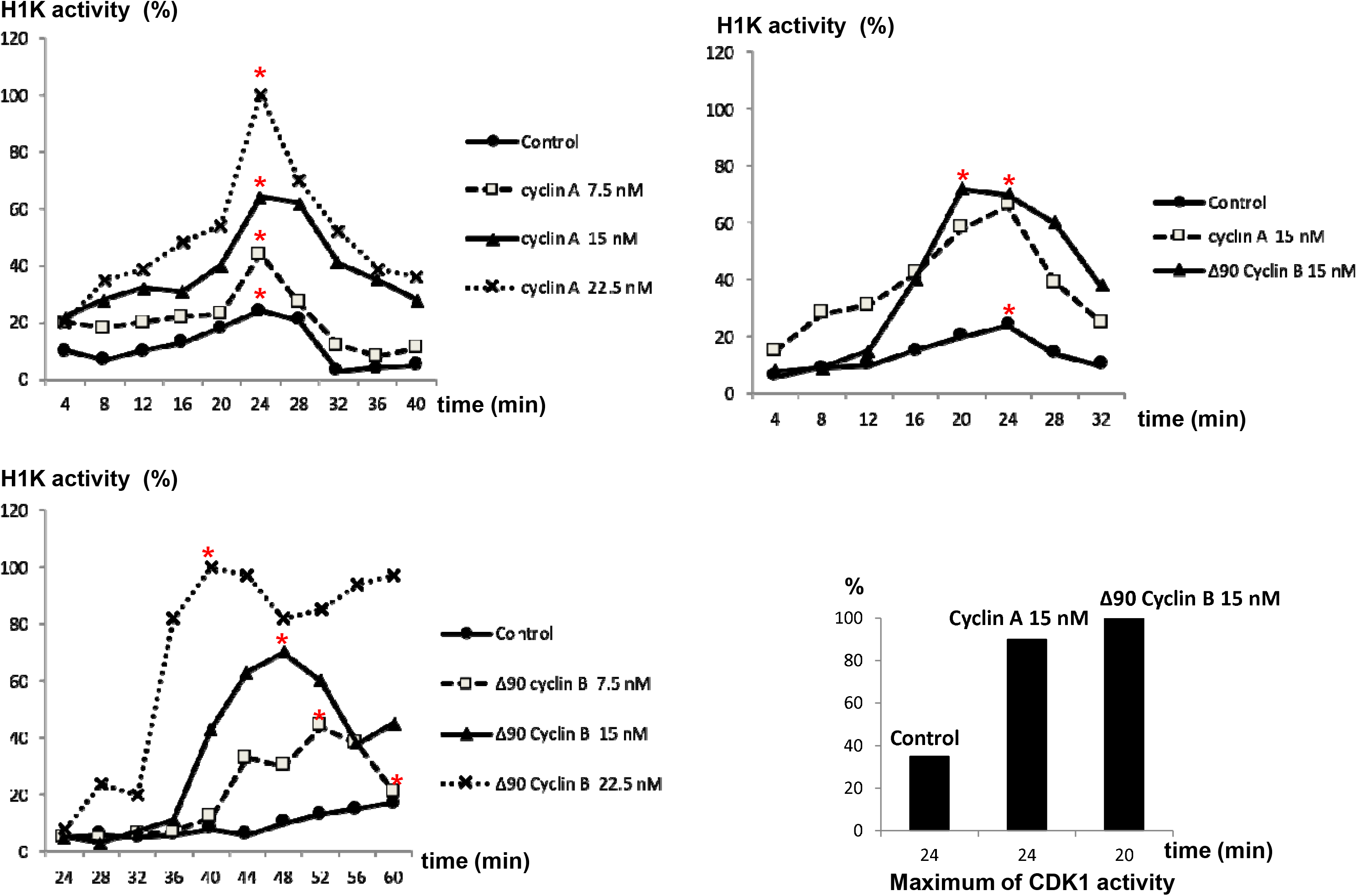
Dose-dependent effects of cyclin A and Δ90 cyclin B addition on the M-phase entry and progression. (A) The cytoplasmic extract was incubated at 21□C in the presence of three concentrations of purified cyclin A or Δ90 cyclin B (7.5, 15, and 22.5 nM) then sampled every 4 min. Samples were analyzed by 8% SDS-PAGE followed by CDC27 Western blot. The dynamics of CDC27 phosphorylation were compared to control. Red asterisks indicate the time points when a substantial shift in CDC27 migration was observed, and which were assumed to be the M-phase entry. Note that the Western blots shown here are the compilation of two separate membranes in each lane because the samples of 4-32 min time points were run and blotted separately from the 36–40 min samples in the series with cyclin A, and the samples of 24-32 min were run and blotted separately from the 36–60 min samples in the series with Δ90 cyclin B. Note different time points of incubation for each experiment (cyclin A *vs*. Δ90 cyclin B) because the timing of M-phase differed from one lot of emryos to the other. (B) The histone H1 kinase assays of the control extract, and the extracts containing increased concentrations of purified cyclin A or Δ90 cyclin B (7.5, 15 and 22.5 nM) corresponding to the CDC27 Western blot shown in Fig. 2A for. Asterisks mark the peaks of histone H1 kinase activity. (C) The histone H1 kinase assays were performed with samples from the same cell-free extract supplemented with purified cyclin A or Δ90 cyclin B (15 nM each) to avoid differences in M-phase timing in different extracts as in Fig. 2A and B. The top panel shows the progression of histone H1 kinase activity in each experimental variant. Asterisks mark the peaks of histone H1 kinase activity. The bottom panel shows histograms depicting the maximum of histone H1 kinase activity obtained in a single reaction series.

In figure 2A (top panel), in the control extract, the up-shift of CDC27 (caused by phosphorylation by the active CDK1) appeared at 24 min of the incubation, and the down-shift, corresponding to the dephosphorylation and thus inactivation of CDK1, at 40 min of incubation. 7.5 nM purified cyclin A does not change the timing of CDC27 phosphorylation during the whole M-phase duration, which remains exactly as in the control. In the presence of 15 nM cyclin A, CDC27 becomes phosphorylated, which marks the M-phase entry, at the same time as in the control, i.e. at 24 min of incubation. However, at 40 min of incubation, there was no downshift of the CDC27 band, which suggests the arrest of the M-phase progression. Interestingly, the addition of 22 nM cyclin A to the extract accelerates CDC27 phosphorylation by 4 min then arrests the M-phase progression similar to the 15 nM cyclin A.

As shown in figure 2A (bottom panel), the timing of the initial up-shift (phosphorylation) of CDC27 accelerated in a dose-dependent manner with the increasing concentration of Δ90 cyclin B in the extracts. The up-shifted band of phosphorylated CDC27 (P-CDC27) appears after 44, 40, and 36 min when Δ90 cyclin B is added to the extracts in 7.5; 15 and 22.5 nM concentration, respectively. This shows that supplementation with exogenous cyclin A and B acts differently on the timing of the initial CDC27 phosphorylation, which, indirectly, show the effect on the timing of CDK1 activation.

To investigate directly how the exogenous cyclin A or cyclin B modifies CDK1 activity in the extract, and how they impact the mitotic progression, we measured the histone H1 kinase activity under the conditions of increasing concentrations of cyclin A or Δ90 cyclin B added to the extracts before the M-phase (Fig. 2B). We found that the concentration of the added cyclin A or B positively correlated with the increase in the activity of H1 kinase. However, the dynamics of histone H1 kinase activity were clearly different for each added cyclin as already suggested by the timing of CDC27 phosphorylation (shown above in figure 2A). The increasing cyclin A-concentration in the extract paralleled the increase in the mitotic H1 kinase activity (Fig. 2B, top panel). Interestingly, in all cyclin A concentrations, the peaks of H1 kinase activation, which mark the peaks of CDK1 activity during the M-phase, were always observed at 24 min of incubation (Fig. 2B, top panel). This indicates that the timing of biochemical events was not modified despite significant differences in the levels of histone H1 kinase activities at different concentrations of the cyclin A. On the other hand, in the Δ90 cyclin B addition experiment, the increase of H1 kinase activity was gradual paralleling the increasing concentration of cyclin B. The fastest increase in histone H1 kinase activation was observed for the highest concentration of added cyclin B (i.e. 22,5 nM). This indicates the acceleration of biochemical events by the increased cyclin B level. Importantly, the time points of the peaks of H1 kinase activity are positively correlated with cyclin B concentrations in the extracts and accordingly shifted in time. The CDK1 activity reached the maximum the soonest in the extract with the maximal dose of cyclin B. Thus, the timing of mitotic events in the extracts is clearly regulated by cyclin B in a dose-dependent manner (Fig. 2B, bottom panel). These differences show that only the increased levels of cyclin B, but not that of cyclin A, accelerate M-phase progression and clearly show different roles of A- and B-type of cyclins in the regulation of the timing of mitotic events.

The results presented in figures 2A and 2B were obtained from two different extracts (note different time points of incubation foe each experiment), therefore some differences between two independent experiments are possible. For this reason, Figure 2C compares the effect of adding 15 nM cyclin A or 15 nM Δ90 cyclin B to the same control extract. This experiment clearly shows the gradual increase of H1 kinase activity upon cyclin A increase (as in the previous experiment shown in fig. 2B, top panel), and the slower initial dynamics of H1 kinase activation followed by a rapid acceleration (as in the previous experiment shown in figure 2B, bottom panel) upon cyclin B increase. Moreover, the amplitudes of CDK1 activity are higher in the extracts supplemented with 15 nM cyclin A (Fig. 2c, top panel) than in the control extract, and become even higher in the extract with 15 nM Δ90 cyclin B (Fig. 2C, top panel). The peaks of CDK1 activity occur at the same time (24 min) in the control and cyclin A-supplemented extracts, while in the extracts with the cyclin B the peak is accelerated by 4 minutes (20 min time point, Fig. 2C, top panel). The H1 kinase activity measurements performed on the same gel and in a single assay confirm the differences in CDK1 activity (shown in the top panel of Fig.2C). These results demonstrate that the exogenous cyclin A and cyclin B have clearly distinct effects on the mitotic extract, and thus differentially regulate M-phase entry and progression. While cyclin B accelerates the peak of CDK1 activity and increases the amplitude of CDK1 activity, cyclin A only increases the amplitude of CDK1 activity without modifying the timing of CDK1 activation.

#### CDC6 regulates the timing of M-phase progression through the endogenous cyclin B and not cyclin A

Previously, we have shown that CDC6, acting as CDK1 inhibitor, regulates the timing and amplitude of CDK1 activity during the M-phase (El Dika et al., 2014a). However, we did not distinguish whether CDC6 inhibits CDK1/cyclin A, CDK1/cyclin B, or both. To distinguish between these possibilities, we followed the timing of M-phase progression in the extract depleted of cyclin A and supplemented with CDC6. We wanted to check whether CDC6 acts differently in the presence of the endogenous cyclin B only (cyclin A-depleted extract) and when both cyclins are present We knew from previously reported data that cyclin A depletion delays the M-phase entry in the extract (Vigneron et al., 2018). To this end, we supplemented the cell-free extract with the pre-immune or immune cyclin A serum at 1:100 dilutions. The pre-immune serum-treated extract served as a control and the immune cyclin A serum depleted cyclin A from the extract (Fig. 3A). The Fig. 3B shows that the addition of 5 nM GST-CDC6 to the control extract delayed M-phase entry (from 16 min time point in the control to 24 min in the presence of exogenous CDC6; Fig. 3B). The same effect was already described by us previously in the paper by El Dika and collaborators (2014a). Next, we compared the effect of 5 nM GST-CDC6 addition to the extract depleted of cyclin A. The experiment showed, as expected from previously published data (Vigneron et al., 2018), that cyclin A depletion delayed the M-phase entry. However, the addition of GST-CDC6 to cyclin A-depleted extract did not change the time course of the M-phase (Fig. 3B, compare the third and the fourth row). These data show that the time regulation of the M-phase by CDC6 is exercised only through CDK1/cyclin B, but not through the CDK1/cyclin A.

**Fig.3.**
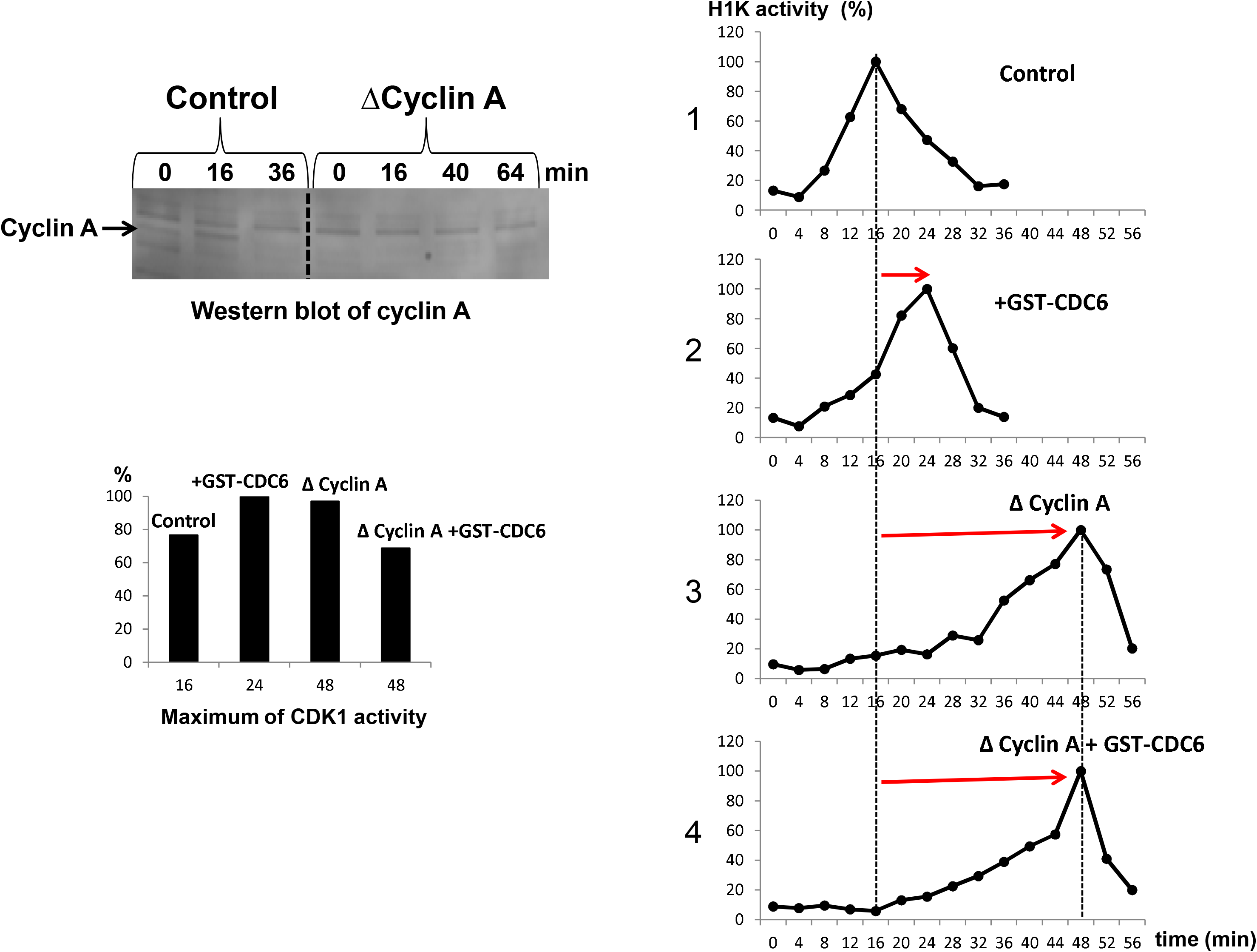
CDC6 regulates M-phase entry timing through the endogenous cyclin B and not through cyclin A. (A) Depletion of cyclin A from the extract was followed by Western blot of *Xenopus laevis* cyclin A. Note the accumulation of cyclin A between 0 and 16min of incubation, and its degradation at 36min of incubation in the control extract (left part of the membrane), and the absence of cyclin A in the depleted extract during 64 min of incubation (right part of the membrane). (B) Quantification of histone H1 kinase activity in control, non-depleted extract (1; top), non-depleted extract supplemented with GST-CDC6 (2; second row), cyclin A-depleted extract (3; Δ cyclin A; third row), and cyclin A-depleted extract supplemented with GST-CDC6 (4; bottom. Red horizontal arrows point to the delay in the timing of the peak of histone H1 kinase activity in the experimental extracts (supplemented with CDC6 and/or depleted of cyclin A) in comparison to the timing in the control extract (the vertical dotted line). (C) The comparison of the values of the peaks of histone H1 kinase activity obtained in a single series of histone H1 kinase assay reaction.

To more accurately analyze the relationship between the levels of histone H1 kinase activity we measured the respective maximums of the activity in a single histone H1 kinase reaction. The highest histone H1 kinase activity (designated as 100%) in this experiment was observed in the presence of GST-CDC6 at the 24 min time point of incubation (Fig. 3C). The maximum peak of H1 kinase activity in the control extract took place at 16 min time point and was lower than in the presence of added GST-CDC6. At first glance, it looks paradoxical because CDC6 inhibits CDK1 activity and one could expect lower histone H1 kinase activity in the GST-CDC6-supplemented extract. We explain this apparent paradox by the concomitant effect of CDC6 on CDK1 activity and the timing of the biochemical events. As the peak of CDK1 activity appeared 8 minutes later in the GST-CDC6-supplemented extract than in the control, more endogenous cyclin B was accumulated in this extract and this resulted in the observed difference in the amplitude of CDK1 activity. The threshold necessary to attain the maximal CDK1 activity is different in each case and results from the interplay between the inhibitory action of exogenous CDC6 and the stimulating action of newly synthetized and accumulated endogenous cyclin B. The maximum activity of CDK1 in the cyclin A-depleted (Δ cyclin A) extract was higher than in the extract supplemented with 5 nM GST-CDC6 (Δ cyclin A+GST-CDC6; Fig. 3C). In these two extracts, the peak of H1 kinase activity appeared at the same time at 48 min of incubation, which suggests that they had the same level of accumulated endogenous cyclin B. Thus, the addition of exogenous CDC6 to the Δ cyclin A extract diminished the level of CDK1 activity because it acted on the equal pool of endogenous cyclin B in each case as it happens upon the addition of CDC6 to the control extracts containing both cyclin A and B (El Dika et al., 2014a).

#### CDC6 is stably associated with CDK1/2 during whole M-phase, while Xic1 dissociates from CDK1/2 complexes specifically during the peak of CDK1 activity

Xic1 is a *bona fide* CDK inhibitor. We wanted to check whether Xic1 associates with the CDK mitotic complexes by precipitating all CDKs by Sepharose p9 beads and analyze whether Xic1 is associated with these complexes, as we did before for a proteomic analysis of CDK complexes in Xenopus laevis embryos (Marteil et al., 2012). To do so, we first determined the timing of the M-phase in the cell-free extract by histone H1 kinase assay to find the critical time points during the M-phase from which the samples for p9 precipitation would be tested. Additionally, Western blotting analysis of the marker proteins CDC27, MCM4, cyclin A1, and cyclin B2 was performed. Subsequently, the selected samples with known mitotic status were used for p9 precipitation followed by a Western blotting analysis of cyclin A1, cyclin B2, CDC6, CDK1, CDK2, and Xic1 on the same blot.

The top part of the fig 4A shows the progression of histone H1 kinase activity during the M-phase in the extract. It starts with a typical slow increase in histone H1 kinase activity until 20 min time point. Then, it reached the maximum activity peak at 24 min of incubation, followed by an immediate drop of CDK1 activity reaching the minimum activity at 32 min of incubation. This dynamic was corrborated by CDC27 and MCM4 phosphorylation state, reflected by the electrophoretic mobility shifts, visualized by Western blotting of the same extract. The maximum up-shift of CDK1-phosphorylated CDC27 and MCM4 was observed between 20 and 24 min of incubation. After 32 min of incubation, the CDC27 and MCM were dephosphorylated (downshifted bands).

**Fig.4.**
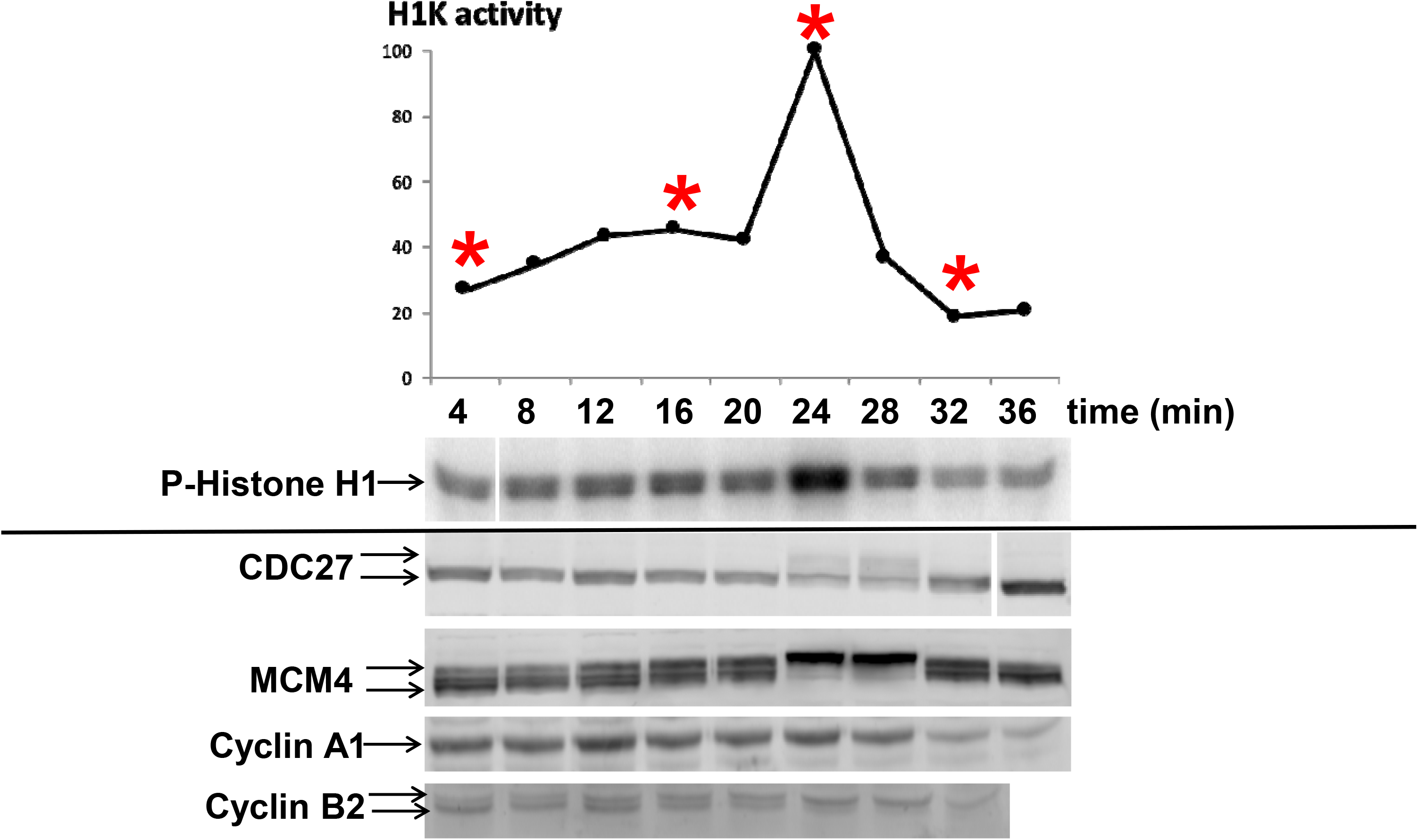
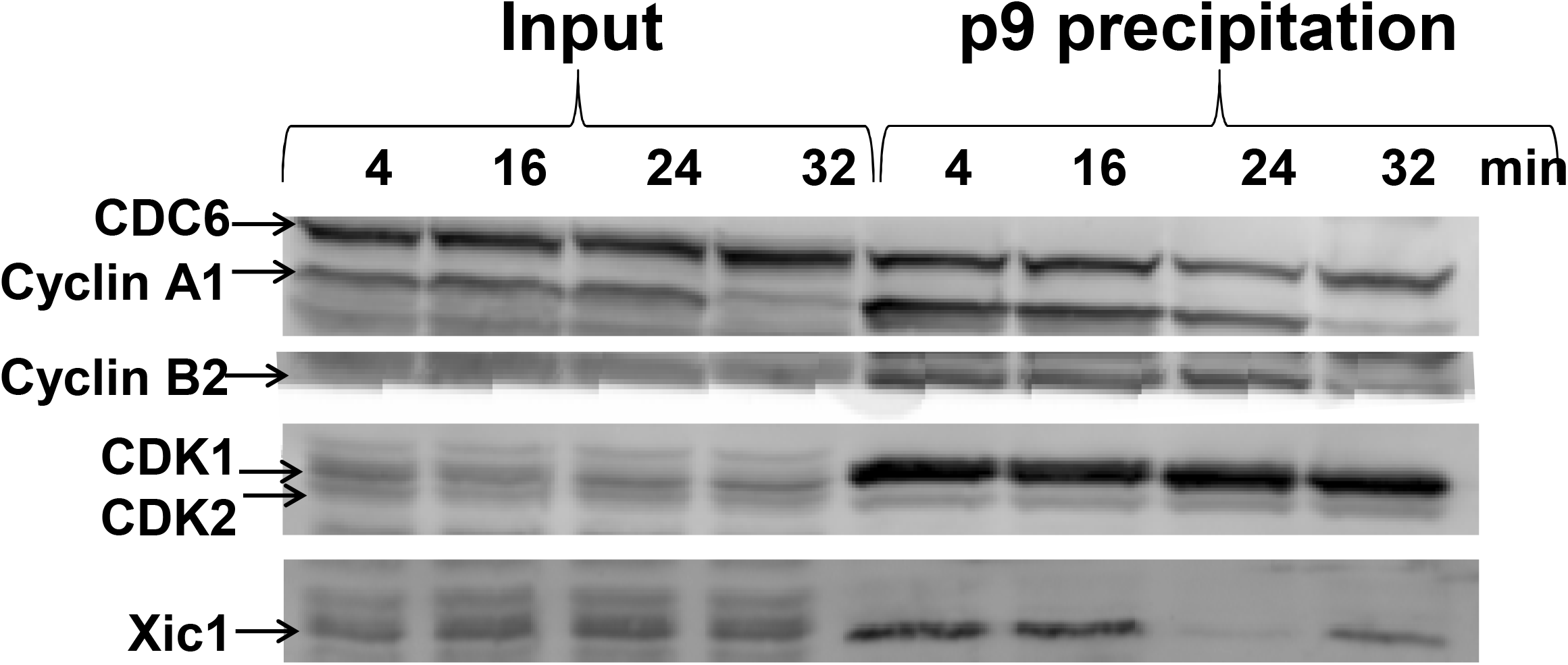
CDC6 and Xic1 associate with p9-precipitated CDK complexes containing also cyclin A and cyclin B. (A) The cytoplasmic extract was incubated at 21□ and sampled every 4 min for histone H1 kinase assay and Western blot analysis. The same samples were split for the histone H1 kinase activity assay, and for 8% SDS-PAGE, followed by CDC27, MCM4, cyclin A1, and cyclin B2 Western blotting. Red asterisks indicate the time points selected based on the basis of histone H1 kinase activity for further Sepharose p9 beads precipitation of CDK complexes during the M-phase. P-histone H1 kinase panel shows the results of the autoradiography assay. Technical remark: note that the first 4 min slot is separated from the following ones because it was revealed on another membrane and two membranes were combined for better visualization. CDC27 and MCM4 Western blots show the M-phase-typical shift of the bands of these proteins due to CDK1 phosphorylation. Technical remark: note that the last 36 min slot of CDC27 is separated from the preceding ones because they were blotted on two different membranes and combined for visualization. Cyclin A1 and cyclin B2 Western blots made from the same samples show accumulation and degradation during the M-phase progression in the extract. (B) Samples of 10 μl of extracts were added to 10 μl p9 beads pre-equilibrated with a buffer supplemented with 1% BSA and protease inhibitors, and p9-associated proteins were precipitated. Samples were analyzed by Western blot with anti-XlCDC6, anti-cyclin A1, anti-cyclin B2, anti-PSTAIR antibody (which detects both CDK1 and CDK2), and anti-Xic1 antibody. Input is shown on the left 4-32 min slots, p9-precipitated complexes are shown on the right 4-32 min slots. In the p9-precipitated sample note the stable levels of CDK1, CDK2, and CDC6, the degradation of cyclin A1 and B2 during the M-phase. Importantly, the drastic reduction of Xic1 is visible specifically at the 24 min time point, while the quantity of p27Xic1 is stable in the input. This indicates that the disappearance of the Xic1 at 24 min time point in p9-precipitated material, and its reappearance at 32 min time point is not due to degradation, but to the dissociation from the CDK complexes.

We also assessed the dynamics of cyclin A1 and cyclin B2 in the extract. Fig 4A (the bottom part) shows that cyclin A1 accumulates gradually, and strongly decreases due to degradation at 32 min of incubation. Cyclin B2 also accumulates gradually and is also gradually changing its phosphorylation status, visible as a gradual up-shift up to 28 min of incubation, followed by degradation at 32 min time point. These results allowed us to choose extract samples for further analysis by Sepharose p9 precipitation and Western blotting.

Fig. 4B shows the comparison of Western blotting of the input extract (left) and the p9 precipitation (right). It shows that CDC6 is stably associated with the mitotic CDKs. Cyclin A1 degradation occurs at the same time in the input and in the p9-precipitate (32 min). Cyclin B2 behaves in the same way. The levels of CDK1 and CDK2 in the input and p9 precipitated material remain stable during the whole M-phase period between 4 and 32 minutes of incubation. However, Xic1, which is stably present during the whole M-phase in the input is clearly absent from the p9 precipitate at 24 min, and rapidly re-associates with CDK complexes at 32 min time point. This dynamic shows that while CDC6 remains stably associated with CDK complexes, the Xic1 dissociates specifically at the time when CDK1 reaches the maximum activity.

### Discussion

We showed here that the cyclin A and cyclin B differentially regulate the timing and progression of the M-phase. The increase in the cyclin A levels positively correlates with the increase in the CDK1 activity but does not change the timing of CDK1 activation or inactivation. This is in contrast to the increase in the cyclin B levels, which although also positively correlate with the increase of CDK1 activity, but contrary to the cyclin A increase accelerates the maximum peak of CDK1 activation. We also showed that the CDC6-dependent lowering of the CDK1 activity during the M-phase, and delaying the dynamics of CDK1 activation acts through the cyclin B, and not through the cyclin A. These results suggests that the CDC6 associates with the CDK1/cyclin B and not with the CDK1/cyclin A and exerts its inhibitory action only on CDK1/cyclin B and not CDK1/cyclin A. Finally, we demonstrated that the *bona fide* CDK inhibitor Xic1 is a part of the mitotic CDK complex and that Xic1 association with CDK is strictly cell cycle-regulated because the Xic1 dissociates from CDK specifically during the mitotic peak of CDK1 activity.

A simple mechanism regulating the timing of mitotic divisions based on the accumulation and degradation of cyclin was proposed already in 1989 (Murray et al., 1989; Murray and Kirschner, 1989). However, this mechanism did not address the precision of the temporal control of the cell division in the early embryo. Cyclin B accumulation to a threshold level is necessary to trigger the M-phase entry. However, as we previously showed, the CDC6-dependent mechanism controls in parallel the timing of the mitotic entry through the regulation of the dynamics of CDK1 activation (El Dika et al., 2014a). CDC6 allows setting the correct timing of M-phase entry, and controls the level of CDK1 activity in cooperation with the continuously increasing level of cyclin B. It is also responsible for the correct shape of the curve of CDK1 activity during the entry into the M-phase, which has a character of a diauxic curve with typical inflection points (Dębowski et al., 2019). Until now, it was not known whether these functions of CDC6 are fulfilled via CDK1/cyclin B or/and CDK1/cyclin A complexes. Our current results demonstrate that this CDC6-dependent control of the mitotic events is exercised only through the cyclin B, and not through cyclin A-containing CDK complexes.

This conclusion creates an apparent paradox because it is known (e.g. Vigneron et al., 2018), and also confirmed by our presented here study that the depletion of endogenous cyclin A strongly delays mitosis. Other studies also confirmed the importance of cyclin A in triggering mitosis entry by showing that CDK1/cyclin A phosphorylates Bora to promote Aurora A-dependent Plk1 phosphorylation, which triggers the activation of MPF amplification loop, and mitotic entry (Vigneron et al., 2018). Thus, cyclin A is indeed involved in the regulation of the dynamic of mitotic entry and progression, however, we demonstrated in the current paper that this control is CDC6-independent.

The CDC6 is not a *bona fide* CDK inhibitor. Moreover, it acts as an activator of CDK2 during the S-phase (Kan et al., 2008a, 2008b; Uranbileg et al., 2012). This happens through the separation of CDK2 form p27. The *bona fide* inhibitor of CDK is the Xic1, a Xenopus p21cip1/p27kip1 family member, identified in *Xenopus laevis* embryos (Su et al., 1995; Finkielstein et al., 2001; Zhongsheng et al., 2002). We demonstrated here that in the Xenopus embryo cycling extracts the CDC6 and Xic1 are in the same complexes. This strengthens the hypothesis that CDC6 may serve as a platform for the assembly of CDK1 and Xic1 to promote CDK1 inhibition. During the S-phase, CDK2 kinase activity is regulated by CDC6 indirectly (Kan et al., 2008a, 2008b; Uranbileg et al., 2012). CDC6 activates CDK2 in somatic cells by the removal of the CDK inhibitor p27 from the CDK2, triggering the kinase activation necessary for the initiation of the S-phase. Our previous studies demonstrated that, during mitosis, the CDC6 has an opposite effect on CDK1 (El Dika et al., 2014a). In this case, CDC6 regulates CDK1 by targeting the CKI inhibitors to the CDK1. A recent study detailing the association of CDK1 with cyclin B, CDC6, and p27 (Cks1) homologs in yeasts (Örd et al., 2019) has shown that, indeed, the CDC6 behaves like a platform gathering all the components of the CDK1 complex. Moreover, these studies showed that the association of these components is coordinated by phosphorylation and dephosphorylation of the LxF motif of CDC6, which confirms the multifunctional regulatory role of CDC6 during M-phase. Our data suggest that in *Xenopus laevis* cell-free extract this regulation may be very similar that in yeast, and that Xic1 may play the role of yeast Cks1. Our observation that Xic1 dissociates from CDK complex specifically when the CDK1 activity level achieves the maximum, suggests that CDC6 and Xic1 actively participate in the rapid activation of CDK1 right before it reaches the maximum. We postulate that, in *Xenopus laevis* cell-free extracts, the rapid and transitional release of Xic1 from the mitotic CDK complex allows the achievement of the extremely steep curve of the CDK1 activity in the M-phase. Our model allows us to unify the role of CKI in the M-phase regulation in lower Eukaryot, the yeast, and the vertebrate *Xenopus laevis.* It seems therefore that in both cases CDK1 is actively inhibited during the M-phase by the CDC6-dependent mechanism including CKI, the Xic1 in the case of Xenopus laevis.

## Acknowledgements

We thank Anna Philpott (University of Cambridge, UK), Marie-Anne Felix (IJM, Paris, France), Marcel Méchali (IGH, Montpellier, France), Daniel Fisher (IGMM, Montpellier, France), Thierry Lorca (CRBM, Montpellier, France), Laurent Meijer and Olivier Lozach (Marine Station, Roscoff, France) for sharing with us antibodies, recombinant proteins and p9 beads. JZK was supported by the grant “Kościuszko” # 5508/2017/DA from the Polish Ministry of National Defense.

## References

Arellano, M., Moreno, S., 1997. Regulation of CDK/cyclin complexes during the cell cycle. Int. J. Biochem. Cell Biol. https://doi.org/10.1016/S1357-2725(96)00178-1

Blain, S.W., Montalvo, E., Massagué, J., 1997. Differential interaction of the cyclin-dependent kinase (CDK) inhibitor p27(Kip1) with cyclin A-Cdk2 and cyclin D2-Cdk4. J. Biol. Chem. 272, 25863–25872. https://doi.org/10.1074/jbc.272.41.25863

Borsuk, E., Jachowicz, J., Kloc, M., Tassan, J.P., Kubiak, J.Z., 2017. Role of Cdc6 during oogenesis and early embryo development in mouse and Xenopus laevis, in: Results and Problems in Cell Differentiation. Springer Verlag, pp. 201–211. https://doi.org/10.1007/978-3-319-44820-6_7

Calzada, A., Sacristán, M., Sánchez, E., Bueno, A., 2001. Cdc6 cooperates with Sic1 and Hct1 to inactivate mitotic cyclin-dependent kinases. Nature 412, 355–358. https://doi.org/10.1038/35085610

Chen, J., Saha, P., Kornbluth, S., Dynlacht, B.D., Dutta, A., 1996. Cyclin-binding motifs are essential for the function of p21CIP1. Mol. Cell. Biol. https://doi.org/10.1128/mcb.16.9.4673

D’Angiolella, V., Costanzo, V., Gottesman, M.E., Avvedimento, E. V., Gautier, J., Grieco, D., 2001. Role for cyclin-dependent kinase 2 in mitosis exit. Curr. Biol. 11, 1221–1226. https://doi.org/10.1016/S0960-9822(01)00352-9

Dębowski M., Szymańska Z., Kubiak JZ., Lachowicz M. (2019). Mathematical Model Explaining the Role of CDC6 in the Diauxic Growth of CDK1 Activity During the M-Phase of the Cell Cycle. Cells 8(12):1537.

Duderstadt, K.E., Berger, J.M., 2008. AAA+ ATPases in the initiation of DNA replication. Crit. Rev. Biochem. Mol. Biol. https://doi.org/10.1080/10409230802058296

El Dika, M., Laskowska-Kaszub, K., Koryto, M., Dudka, D., Prigent, C., Tassan, J.P., Kloc, M., Polanski, Z., Borsuk, E., Kubiak, J.Z., 2014a. CDC6 controls dynamics of the first embryonic M-phase entry and progression via CDK1 inhibition. Dev. Biol. 396, 67–80. https://doi.org/10.1016/j.ydbio.2014.09.023

El Dika, M., Dudka, D., Prigent, C., Tassan, J.P., Kloc, M., Kubiak, J.Z., 2014b. Control of timing of embryonic M-phase entry and exit is differentially sensitive to CDK1 and PP2A balance. Int. J. Dev. Biol. 58, 767–774. https://doi.org/10.1387/ijdb.140101jk

Elsasser, S., Lou, F., Wang, B., Campbell, J.L., Jong, A., 1996. Interaction between yeast Cdc6 protein and B-type cyclin/Cdc28 kinases. Mol. Biol. Cell 7, 1723–1735. https://doi.org/10.1091/mbc.7.11.1723

Evans, T., Rosenthal, E.T., Youngblom, J., Distel, D., Hunt, T., 1983. Cyclin: A protein specified by maternal mRNA in sea urchin eggs that is destroyed at each cleavage division. Cell 33, 389–396. https://doi.org/10.1016/0092-8674(83)90420-8

Finkielstein CV, Lewellyn AL, Maller JL. (2001). The midblastula transition in Xenopus embryos activates multiple pathways to prevent apoptosis in response to DNA damage. Proc Natl Acad Sci U S A. 98(3): 1006–1011.

Girard, F., Strausfeld, U., Fernandez, A., Lamb, N.J.C., 1991. Cyclin a is required for the onset of DNA replication in mammalian fibroblasts. Cell 67, 1169–1179. https://doi.org/10.1016/0092-8674(91)90293-8

Glotzer, M., Murray, A.W., Kirschner, M.W., 1991. Cyclin is degraded by the ubiquitin pathway. Nature 349, 132–138. https://doi.org/10.1038/349132a0

Gopinathan, L., Ratnacaram, C.K., Kaldis, P., 2011. Established and novel Cdk/Cyclin complexes regulating the cell cycle and development. Results Probl. Cell Differ. 53, 365–389. https://doi.org/10.1007/978-3-642-19065-0_16

Huang, J.Y., Morley, G., Li, D., Whitaker, M., 2007. Cdk1 phosphorylation sites on Cdc27 are required for correct chromosomal localisation and APC/C function in syncytial Drosophila embryos. J. Cell Sci. 120, 1990–1997. https://doi.org/10.1242/jcs.006833

Kan, Q., Jinno, S., Kobayashi, K., Yamamoto, H., Okayama, H., 2008a. Cdc6 determines utilization of p21WAF1/CIP1-dependent damage checkpoint in S phase cells. J. Biol. Chem. 283, 17864–17872. https://doi.org/10.1074/jbc.M802055200

Kan, Q., Jinno, S., Yamamoto, H., Kobayashi, K., Okayama, H., 2008b. ATP-dependent activation of p21WAF1/CIP1-associated Cdk2 by Cdc6. Proc. Natl. Acad. Sci. U. S. A. 105, 4757–4762. https://doi.org/10.1073/pnas.0706392105

King, R.W., Jackson, P.K., Kirschner, M.W., 1994. Mitosis in transition. Cell 79, 563–571. https://doi.org/10.1016/0092-8674(94)90542-8

Lorca, T., Devault, A., Colas, P., Van Loon, A., Fesquet, D., Lazaro, J.B., Dorée, M., 1992. Cyclin A-Cys41 does not undergo cell cycle-dependent degradation in Xenopus extracts. FEBS Lett. 306, 90–93. https://doi.org/10.1016/0014-5793(92)80844-7

Marteil, G., Gagné, J.P., Borsuk, E., Richard-Parpaillon, L., Poirier, G.G., Kubiak, J.Z., 2012. Proteomics reveals a switch in CDK1-associated proteins upon M-phase exit during the Xenopus laevis oocyte to embryo transition. Int. J. Biochem. Cell Biol. 44, 53–64. https://doi.org/10.1016/j.biocel.2011.09.003

Müller-Tidow, C., Ji, P., Diederichs, S., Potratz, J., Bäumer, N., Köhler, G., Cauvet, T., Choudary, C., van der Meer, T., Chan, W.-Y.I., Nieduszynski, C., Colledge, W.H., Carrington, M., Koeffler, H.P., Restle, A., Wiesmüller, L., Sobczak-Thépot, J., Berdel, W.E., Serve, H., 2004. The Cyclin A1-CDK2 Complex Regulates DNA Double-Strand Break Repair. Mol. Cell. Biol. 24, 8917–8928. https://doi.org/10.1128/mcb.24.20.8917-8928.2004

Murray, A.W., Kirschner, M.W., 1989. Cyclin synthesis drives the early embryonic cell cycle. Nature 339, 275–280. https://doi.org/10.1038/339275a0

Murray, A.W., Solomon, M.J., Kirschner, M.W., 1989. The role of cyclin synthesis and degradation in the control of maturation promoting factor activity. Nature 339, 280–286. https://doi.org/10.1038/339280a0

Örd M, Venta R, Möll K, Valk E, Loog M. (2019). Cyclin-Specific Docking Mechanisms Reveal the Complexity of M-CDK Function in the Cell Cycle. Mol Cell. 11; 75(1):76–89.e3.

Pagano, M., Pepperkok, R., Verde, F., Ansorge, W., Draetta, G., 1992. Cyclin A is required at two points in the human cell cycle. EMBO J. 11, 961–971. https://doi.org/10.1002/j.1460-2075.1992.tb05135.x

Petersen, B.O., Wagener, C., Marinoni, F., Kramer, E.R., Melixetian, M., Denchi, E.L., Gieffers, C., Matteucci, C., Peters, J.M., Helin, K., 2000. Cell cycle- and cell growth-regulated proteolysis of mammalian CDC6 is dependent on APC-CDH1. Genes Dev. 14, 2330–2343. https://doi.org/10.1101/gad.832500

Pines, J., 1995. Cyclins and cyclin-dependent kinases: A biochemical view. Biochem. J. https://doi.org/10.1042/bj3080697

Pines, J., Hunter, T., 1991. Human cyclins A and B1 are differentially located in the cell and undergo cell cycle-dependent nuclear transport. J. Cell Biol. 115, 1–17. https://doi.org/10.1083/jcb.115.1.1

Rechsteiner, M., Rogers, S.W., 1996. PEST sequences and regulation by proteolysis. Trends Biochem. Sci. https://doi.org/10.1016/S0968-0004(96)10031-1

Russo, A.A., Jeffrey, P.D., Patten, A.K., Massagué, J., Pavletich, N.P., 1996. Crystal structure of the p27(Kip1) cyclin-dependent-kinase inhibitor bound to the cyclin A-Cdk2 complex. Nature 382, 325–331. https://doi.org/10.1038/382325a0

Sherr, C.J., Roberts, J.M., 1999. CDK inhibitors: Positive and negative regulators of G1-phase progression. Genes Dev. https://doi.org/10.1101/gad.13.12.1501

Su JY, Rempel RE, Erikson E, Maller JL (1995). Cloning and characterization of the Xenopus cyclin-dependent kinase inhibitor p27XIC1. Proc Natl Acad Sci U S A. 92(22): 10187–10191.

Uranbileg, B., Yamamoto, H., Park, J.H., Mohanty, A.R., Arakawa-Takeuchi, S., Jinno, S., Okayama, H., 2012. Cdc6 protein activates p27 KIP1-bound Cdk2 protein only after the bound p27 protein undergoes C-terminal phosphorylation. J. Biol. Chem. 287, 6275–6283. https://doi.org/10.1074/jbc.M111.318295

Vigneron, S., Sundermann, L., Labbé, J.C., Pintard, L., Radulescu, O., Castro, A., Lorca, T., 2018. Cyclin A-cdk1-Dependent Phosphorylation of Bora Is the Triggering Factor Promoting Mitotic Entry. Dev. Cell 45, 637–650.e7. https://doi.org/10.1016/j.devcel.2018.05.005

Walker, D.H., Maller, J.L., 1991. Role for cyclin A in the dependence of mitosis on completion of DMA replication. Nature 354, 314–317. https://doi.org/10.1038/354314a0

Yim, H., Erikson, R.L., 2010. Cell division cycle 6, a mitotic substrate of polo-like kinase 1, regulates chromosomal segregation mediated by cyclin-dependent kinase 1 and separase. Proc. Natl. Acad. Sci. U. S. A. 107, 19742–19747. https://doi.org/10.1073/pnas.1013557107

You, Z., Harvey, K., Kong, L., Newport, J., 2002. Xic1 degradation in Xenopus egg extracts is coupled to initiation of DNA replication. Genes Dev. 16, 1182–1194. https://doi.org/10.1101/gad.985302

Zhongsheng You, Kevin Harvey, Lindsay Kong, John Newport (2002). Xic1 degradation in Xenopus egg extracts is coupled to initiation of DNA replication. Genes Dev. 16(10): 1182–1194.

Zhu, L., Harlow, E., Dynlacht, B.D., 1995. p107 uses a p21CIP1-related domain to bind cyclin/cdk2 and regulate interactions with E2F. Genes Dev. 9, 1740–1752. https://doi.org/10.1101/gad.9.14.1740

